# Within-electrode temporal envelope processing predicts multi-channel speech outcomes across cochlear implant pulse rates

**DOI:** 10.64898/2026.06.24.734273

**Authors:** Mahan Azadpour, Jonathan Neukam, Nicole Capach, Mario Svirsky

## Abstract

Cochlear implants (CIs) restore hearing by stimulating auditory neurons to encode amplitude envelopes across frequency bands, providing essential cues for speech recognition. This study investigated how stimulation pulse rate constrains temporal envelope processing and speech cue perception in ten post-lingually deaf CI users by evaluating amplitude modulation (AM) detection thresholds and consonant identification performance across pulse rates. The effects of pulse rate on temporal processing and speech perception were examined using both standard clinical multi-channel strategies and single-channel strategies designed to isolate within-channel envelope representations. Results revealed a significant decline in AM detection and consonant recognition performance at the lowest tested pulse rate of 125 pulses per second (pps), consistent with perceptual constraints on temporal processing at low carrier rates, rather than inadequate envelope sampling. At the highest pulse rate of 4000pps, a non-significant reduction in AM detection was observed which may be consistent with previously reported reductions in amplitude discrimination at high pulse rates. Consonant recognition performance remained stable across clinically relevant pulse rates (250–2000pps), though listener-specific pulse rate effects were observed. Notably, significant correlations were found between single-channel and multi-channel performance in AM detection and consonant recognition tasks. These findings support an important contribution of within-electrode temporal envelope processing to multi-channel speech perception and highlight the clinical relevance of individual variability in pulse rate effects.

## 1. Introduction

Cochlear implant (CI) devices restore hearing by electrically stimulating auditory neurons within the cochlea. The functionality of CIs relies on sound coding strategies that transform environmental sounds into electrical pulses, delivered via intra-cochlear electrodes. These strategies are designed to primarily encode amplitude envelopes across frequency bands, as these envelopes convey critical temporal cues for speech recognition. Typical CI strategies represent amplitude envelopes using constant-rate carrier pulses, often sacrificing fine temporal detail.

CI performance depends on accurate encoding of temporal envelope cues, which are critical for speech perception. The ability to perceive these cues can be hindered by inadequate stimulation parameters or listener-specific auditory processing limitations. The repetition rate of carrier pulses likely plays a key role in determining access to speech temporal envelope information. While low carrier pulse rates are expected to distort envelope modulations and impair speech perception, higher pulse rates should, in principle, enhance temporal modulation precision. Despite these theoretical predictions, studies investigating the impact of pulse rate on speech perception have yielded inconsistent results. Some studies demonstrate benefits from moderate-to-high pulse rates (Kiefer et al., 2001; Loizou et al., 2000; Nie et al., 2006; Verschuur, 2005), and some others report either no discernible advantage across pulse rates (Berg et al., 2023; Friesen et al., 2005; Plant et al., 2007; Shannon et al., 2011) or benefits at lower pulse rates (Brochier et al., 2017; Vandali et al., 2000). Furthermore, several studies have identified individually optimal pulse rates, which vary among CI users with no consistent pattern (Plant et al., 2007; Shader et al., 2020; Weber et al., 2007). These discrepancies demonstrate the complexity of speech temporal envelope transmission and suggest that optimal encoding of envelopes with CIs may depend on individual auditory profiles.

We hypothesized that the mixed benefits from CI stimulation pulse rates are driven by the complex effects of pulse rate on envelope transmission. Specifically, we propose that both low and high pulse rates impair psychophysical sensitivity to amplitude modulations (AM). This hypothesis is supported by animal CI studies showing reduced cortical phase locking to AM at high pulse rates (Middlebrooks, 2008) and human CI studies demonstrating declines in AM detection and amplitude increment discrimination as pulse rate increases (Galvin, Fu, 2005; Galvin, Fu, 2009). However, these declines may not necessarily reflect psychophysical impairments at high pulse rates. Instead, reduced sensitivity to electric amplitude increments may primarily be due to a shallower loudness growth with increasing electric amplitude, rather than auditory processing deficits that directly impact speech perception. While shallower loudness growth diminishes sensitivity to electric amplitude increments (Galvin, Fu, 2009), compensatory acoustic-to-electric mappings in CI sound coding strategies can largely offset these effects, preserving speech recognition performance.

To disentangle pulse rate effects on loudness growth and auditory sensitivity to amplitude variations, a prior study employed a novel psychophysical approach to measure the just-noticeable-difference (JND) in current amplitude for high-rate and low-rate pulse trains (Azadpour et al., 2018). The patterns of current JNDs, analyzed using signal detection theory, revealed greater trial-to-trial variability in perceived loudness for the high-rate pulse train. This greater loudness variability heightens uncertainty in detecting amplitude changes and reduces the number of discriminable intensity steps across the hearing range, potentially impairing perception of acoustic envelopes and speech cues.

This study was designed to directly investigate the effect of pulse rate on sensitivity to acoustic envelopes and on speech perception. To eliminate the influence of loudness growth at different pulse rates, AM sensitivity was assessed acoustically using CI processing strategies that account for differences in electric loudness growth. Speech perception was evaluated using consonant identification, a task heavily reliant on temporal modulation cues. AM detection and consonant identification tasks were conducted via standard clinical multi-channel ACE (Advanced Combination Encoder) strategies as well as single-channel programs. Single-channel programs were employed to eliminate the influence of cross-channel spectral representations by encoding the overall signal envelope using a single CI electrode (Azadpour, McKay, 2014). A wide range of ACE channel rates was tested, with electric threshold and comfortable levels determined individually for each pulse rate. It was hypothesized that the transmission of acoustic envelope cues would decrease at both low and high stimulation pulse rates. It was further hypothesized that single-channel AM detection and speech recognition predict variability in outcomes with clinical multi-channel strategies. Results are interpreted with respect to the role of within-electrode temporal processing in CI speech perception.

## 2. Methods

### 2.1. Participants

Ten post-lingually deafened adult CI users of Cochlear Nucleus devices with perimodiolar electrode arrays participated in this study. Subject’s demographics, implant type and the ACE channel rate in their clinical processor are shown in Table I. All participants had been using their CI for at least one year at the time of testing. Subjects with bilateral CIs were tested with their better ear. Participant recruitment and testing were approved by Institutional Review Board (IRB) and informed consent was obtained from all participants.

**Table I.** Subject demographics and device information.

| Subject | Device | Clinical Rate | Clinical Maxima | Etiology | Duration of Hearing Loss (years) | Duration of CI Use (years) | Clinical CNC Word Scores |
| --- | --- | --- | --- | --- | --- | --- | --- |
| 1 | CI24RE-CA | 900 | 8 | Unknown, gradual progressive | 8 (7 hearing aid use) | 4.8 | 66% at 60 dBA |
| 2 | CI24RE-CA | 720 | 8 | Congenital, gradual progressive | 65 (60 hearing aid use) | 7.5 | 50% at 60 dBA |
| 3 | CI24RE-CA | 900 | 10 | Unknown, sudden | 1 | 9.0 | N/A |
| 4 | CI24RE-CA | 900 | 8 | Congenital, gradual progressive | 45 (40 hearing aid use) | 11.8 | N/A |
| 5 | CI24RE-CA | 900 | 10 | Unknown, gradual progressive | 51 (17 hearing aid use) | 11.0 | 68% at 50 dBHL |
| 6 | CI532 | 900 | 8 | Unknown, gradual progressive | 17 (15 hearing aid use) | 4.3 | 80% at 60 dBA |
| 7 | CI532 | 900 | 8 | Unknown, gradual progressive | 41 (28 hearing aid use) | 3.6 | 90% at 60 dBA |
| 8 | CI24RE-CA | 900 | 8 | Congenital, gradual progressive | 10 (9 hearing aid use) | 16.9 | 62% at 60 dBA |
| 9 | CI612 | 900 | 8 | Meningitis, gradual progressive | 1 | 1.1 | 62% at 60 dBA |
| 10 | CI24RE-CA | 900 | 8 | Congenital, gradual progressive | 39 (2 hearing aid use) | 12.0 | 46% at 60 dBA |

### 2.2. Experimental CI Processing Strategies

Audio stimuli were processed and streamed to subjects’ implants using the CCi-MOBILE research platform, bypassing participants’ clinical processors. CCi-MOBILE is a portable, real-time processing and streaming platform developed by the Center for Robust Speech Systems at the University of Texas at Dallas for Cochlear Corporation devices (Azadpour et al., 2025; Ghosh et al., 2022; Shekar, Hansen, 2021). It receives electrical stimulus data from a computing device (e.g., PC, tablet, or smartphone) and streams to the internal CIC3/CIC4 receiver-stimulator of Cochlear Nucleus devices. The CI sound coding strategies are hosted in the computing device.

Experimental multi-channel and single-channel ACE processing strategies were implemented using the CCi-MOBILE Software Suite in Matlab. The CCi-MOBILE software package uses the Nucleus Toolbox for Matlab (https://github.com/Cochlear-Research/nucleus-toolbox) provided by Cochlear Corporation for emulating the ACE processing strategy in clinical CI processors. The multi-channel ACE strategy programs used the active electrodes in the subject’s clinical processor and five experimental channel rates between 125 and 2000pps: 125, 250, 500, 1000, and 2000pps. The channel rates were powers of 2 multiples of 125 to ensure uniform pulse generation by CCi-MOBILE (Azadpour et al., 2025). The number of maxima was set to 6, to ensure that the overall stimulation rate (i.e. channel rate times maxima selection) was below 14,000pps, the maximum rate supported by CCi-MOBILE. This represented a reduction from some participants’ clinical maxima settings (Table I), which is acknowledged as a potential limitation. The remaining strategy parameters, such as frequency allocation, matched the clinical map that the subjects used daily during the period of participation in the study. Single-channel programs used electrode 12 in the middle of the Nucleus CI electrode array, or the closest usable electrode. The tested channel rates for the single-channel strategies included six rates between 125 and 4000pps: 125, 250, 500, 1000, 2000, and 4000pps.

Threshold (T) and comfortable loudness (C) levels were measured for every other active electrode and the levels for the intermediate electrodes were interpolated. T levels were obtained using the method of adjustment. The experimenter adjusted the current level of a pulse train presented to the target electrode at the test rate to find hearing threshold. C levels were estimated by loudness balancing pulse trains at each test rate to a 1000pps reference pulse train at C level. The C level for 1000pps pulse train was obtained using the method of adjustment. Loudness balancing was performed in an adaptive two-interval two-alternative forced choice task, where the stimuli in the two intervals were the test pulse train and the reference 1000pps pulse train. Subjects’ task was to identify the louder interval. The current level of the test pulse train was adapted in 1-up 1-down procedure to find equal loudness to the 1000pps reference. The initial step size was 4 clinical level (CL) for the first two reversals and was reduced to 2 CL in the next 8 reversals. The average level of the last 6 reversals was used as the loudness balanced level. The duration of pulse trains was 500ms for all procedures.

### 2.3. Experiment 1: Consonant Identification

Consonant identification was evaluated using a closed-set paradigm. The audio stimuli were pre-recorded vowel-consonant-vowel syllables described in previous publications (Shannon et al., 1999). The vowels surrounding the middle consonant were both /a/. The consonant was from a set of 16 consonants: /b/, /d/, /g/, /t/, /k/, /p/, /m/, /n/, /v/, /j/, /f/, /z/, /ch/, /sh/, /s/, /l/. The syllable stimuli were imported from audio wave files, which were normalized to have root mean square (RMS) value equal to 0.1. This was the maximum achievable RMS without clipping. The stimuli were processed by the experimental ACE strategies in a real-time manner and streamed to the subjects’ implant via CCi-MOBILE.

A block of stimuli contained a total of 96 syllables with each of the 16 consonants produced by 6 different speakers (3 female, 3 male). For each strategy condition, three blocks of stimuli were presented (288 syllables total), or two blocks (192 syllables) when testing time was limited. Subjects’ task was to select the syllable they heard from the list of choices on a computer screen. Participants received no training on the task, and feedback on their responses was not provided. The multi-channel conditions were completed before the single-channel conditions and the conditions with different pulse rates were counterbalanced. The results were used to construct consonant confusion matrices, from which percent correct scores and information transfer of articulation features were calculated (Azadpour, McKay, 2014; Azadpour et al., 2014; Miller, Nicely, 1955).

### 2.4. Experiment 2: Amplitude Modulation (AM) Detection

Stimuli for AM detection consisted of sinusoidally amplitude-modulated noise bands processed using experimental ACE strategy conditions that varied in channel rate and number of channels, as described in section 2.3. The carrier was random noise low-pass filtered below 6 kHz with a 6th-order Butterworth filter. The carrier noise bands were regenerated for each presentation at full scale between −1 and 1 and then scaled to prevent clipping after AM. This scaling ensured that 100% AM depth remained within the full-scale range, eliminating the risk of distortion in the AM stimuli. AM stimuli were created by modulating noise-band carriers with 25 Hz sinusoids at random phases. This AM frequency was at least five times smaller than all carrier rates, meeting the criteria for accurate envelope sampling as established by modulation discrimination tasks. Performance in these tasks remained accurate when the carrier rate was at least four times the modulation frequency (McKay et al., 1994). Furthermore, the 25Hz AM frequency represents the frequency range of critical speech envelope cues, which are predominantly below 20Hz (Sinha, Azadpour, 2024; Souza, Rosen, 2009).

The detection threshold for AM depth was measured in an adaptive 3-interval 3-alternative forced choice task. In each trial, two intervals contained non-AM noise, while one interval, presented in random order, contained AM noise. Participants were tasked with identifying the interval containing the AM noise. The duration of the stimulus in each interval was 500ms and the inter-stimulus interval was 800ms. The levels of the noise stimuli in each trial were roved by ±3 dB to eliminate potential loudness cues that might influence participants’ responses in AM detection tasks (McKay, Henshall, 2010). The AM depth was adjusted using a 2-down, 1-up adaptive rule, targeting 71% detection accuracy (Levitt, 1971). The initial AM depth was set to ensure AM stimulus was clearly identified. The step size for AM depth was 4% for the first two reversals and reduced to 2% thereafter. The procedure was terminated after 10 reversals and AM depth threshold was the average depth value from the last six reversals.

### 2.5. Statistical Analysis

Repeated measures ANOVA analyses were used to evaluate the effects of pulse rate on different outcome measures, including T and C levels, AM detection thresholds and consonant recognition performance. Post-hoc comparisons used Holm-Sidak corrections for multiple comparisons. The within-subject effects of pulse rate were analyzed using a conservative within-subject approach, where significant effects were determined based on non-overlapping 95% confidence intervals. The correlations between different outcome measures and between single-channel and multi-channel measures were assessed using Pearson correlation analysis.

## 3. Results

### 3.1. T and C Levels

The T and C levels decreased as the stimulation rate increased, consistent with the findings from previous studies (Kreft et al., 2004; Zhou et al., 2012). Figure 1 shows the average T and C levels (in clinical level units, CL) obtained from the 10 subjects at different electrodes across the array and five pulse rates: 125, 250, 500, 1000, and 2000pps. Error bars represent standard errors.

**Figure 1.**
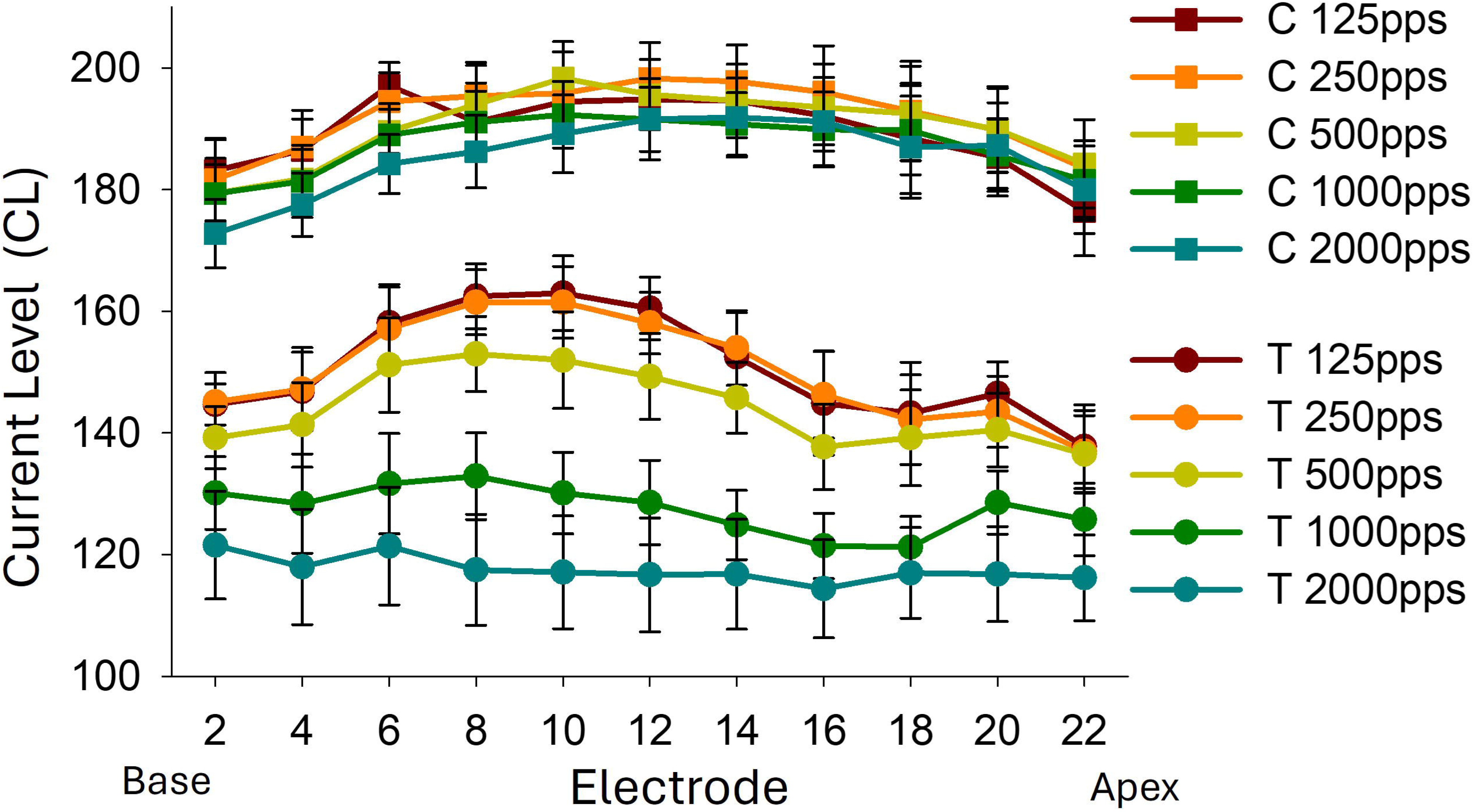
Subject-averaged threshold (T) and comfort (C) levels across pulse rates and electrode locations. Error bars represent standard errors.

The average T level across subjects and electrodes showed a systematic reduction of 33CL (from 151 to 118CL), as pulse rate increased from 125 to 2000pps. A two-way repeated measures ANOVA analysis revealed a significant effect of pulse rate (F(4, 24)=73, p<0.001), no significant effect of electrode location (F(10, 60)=1.34, p=0.23), and a significant interaction between pulse rate and electrode location (F(40, 240)=2.7, p<0.001). Holm-Sidak post-hoc tests showed significant differences between the T levels of mid-array electrodes 8 and 10 and the most apical electrode 22 at 125pps and 250pps only (p<0.05), with no significant differences at higher pulse rates. The average C levels were 189, 192, 190, 187, and 185CL for pulse rates of 125, 250, 500, 1000, and 2000pps, respectively. C levels reduced by 4CL from 125 to 2000pps, and by 7CL from 250 to 2000pps. ANOVA analysis showed significant main effects for pulse rate (F(4, 24)=25.7, p<0.001) and electrode location (F(10, 60)=2.1, p=0.04), as well as a significant interaction (F(40, 240)=1.66, p=0.01). Post-hoc analysis confirmed significant differences between the C levels of electrodes 6 through 16 and the C level of the most apical electrode 22 at 125pps only, with no significant effect of electrode location at higher pulse rates. Analysis of dynamic range (the difference between C and T levels) confirmed a larger impact of pulse rate on T levels than C levels. ANOVA analysis revealed significant effects of pulse rate (F(4, 24)=37.6, p<0.001) and electrode location (F(10, 60)=2.2, p=0.03) on dynamic range, as well as a significant interaction (F(40, 240)=1.78, p=0.005). Dynamic range significantly increased at higher pulse rates, driven by larger reduction in T levels compared to C levels.

### 3.2. Consonant Identification

Closed-set identification results were used to generate consonant confusion matrices for each experimental strategy condition. Percent correct consonant scores were calculated as the ratio of the sum of diagonal elements to the sum of all elements in the confusion matrices. Additionally, patterns of confusion between consonant articulation features were analyzed by grouping consonants based on manner of articulation, place of articulation, and voicing features. Manner of articulation was categorized as plosives/affricates (/b/, /d/, /g/, /t/, /k/, /p/, /j/, /ch/), nasals (/m/, /n/), and fricatives/approximants (/v/, /f/, /z/, /sh/, /s/, /l/). Place of articulation was categorized as front (/b/, /p/, /m/, /v/, /f/), middle (/d/, /t/, /n/, /z/, /s/, /l/), and back (/g/, /k/, /j/, /ch/, /sh/). Voicing categories comprised voiced (/b/, /d/, /g/, /m/, /n/, /v/, /j/, /z/, /l/) and unvoiced (/t/, /k/, /p/, /f/, /ch/, /sh/, /s/). Confusions between these articulation categories were analyzed using information transfer analysis (Miller, Nicely, 1955).

The average correct consonant scores and information transfer of consonant features are presented in Figure 2 and Table II. The top panel of Figure 2 displays average correct consonant scores, while the bottom panels show information transfer of consonant features. Error bars indicate standard errors. Consonant identification performance, as expected, was significantly better for multi-channel ACE strategy conditions (blue squares) compared to single-channel strategy conditions (green circles), indicting the reliance of speech perception on cross-channel spectral information. Nevertheless, consonant information transmission under most single-channel strategy conditions exceeded the chance level and its 95% confidence interval, represented by solid and dashed horizontal lines, respectively. The chance level for correct consonant scores was 6.25%, based on 16 response choices. For information transfer analysis the chance level is 0%. The 95% confidence interval for correct consonant score was calculated using a binomial distribution (Thornton, Raffin, 1978). For information transfer analysis, 95% confidence intervals were derived by simulating chance distributions from random confusion matrices (Azadpour et al., 2014).

**Figure 2.**
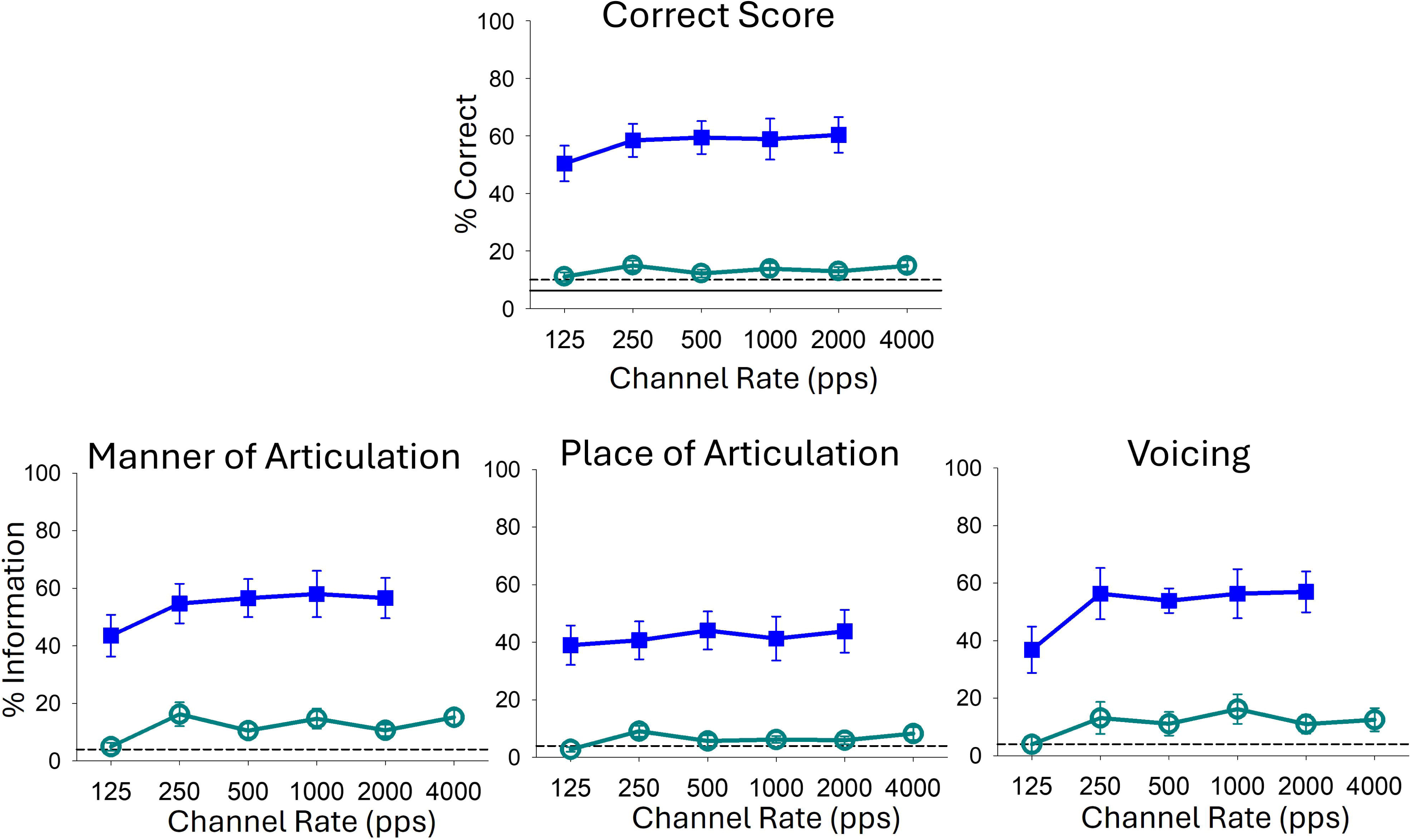
Average consonant identification performance for single-channel (green circles) and multi-channel clinical (blue squares) strategies across pulse rates. The top panel shows overall consonant scores and the bottom panels show information transfer of articulation features. Errors bars represent standard errors.

**Table II.**
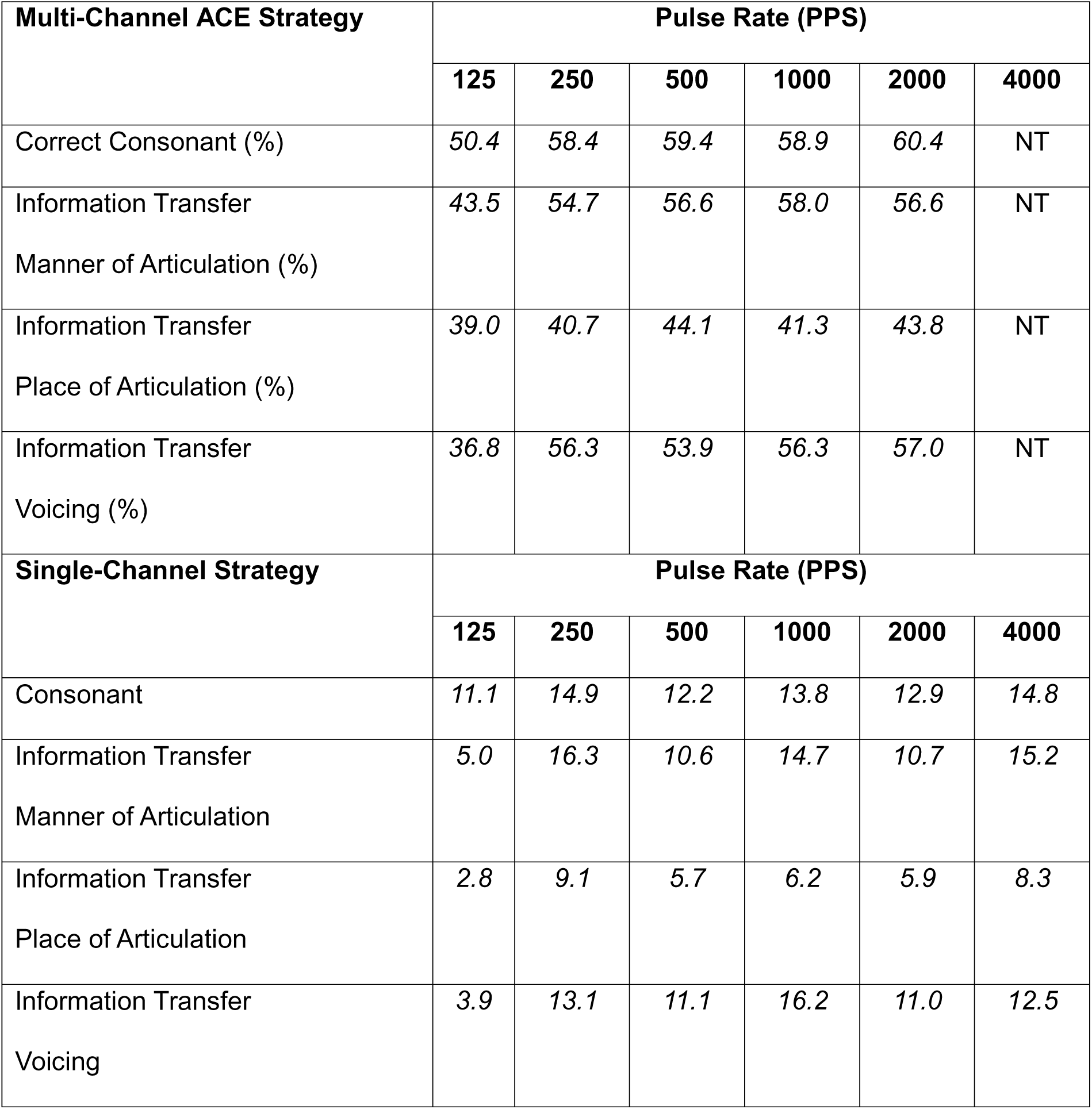
Average consonant identification performance results across pulse rates.

The effects of pulse rate on consonant scores and information transfer of consonant features were assessed using separate one-way repeated measures ANOVA analyses for multi-channel and single-channel strategies. For multi-channel strategies, pulse rate significantly influenced correct consonant scores (F(4,36)=2.8, p=0.042) and information transfer for manner of articulation feature (F(4, 36)=3.4, p=0.021), while no significant effect was observed for place of articulation (F(4, 36)=0.81, p=0.52) or voicing features (F(4, 36)=2.2=0.096). Post-hoc analysis indicated that the significant effects were primarily driven by poorer performance at 125 pps (p < 0.05 vs. all other rates). There was no significant difference between 250, 500, 1000, and 2000 pps conditions (p>0.05).

For single-channel strategies, pulse rate did not have a significant effect on correct consonant score (F(5, 45)=1.37, p=0.26). However, significant effects of pulse rate were observed for information transmission of manner of articulation feature (F(5, 45)=3.3, p=0.013) and place of articulation (F(5, 45)=4.36, p=0.003), with no significant effect on voicing feature (F(5, 45)=2.1, p=0.09). Post hoc analysis revealed that significant effects of pulse rate were driven by poorer performance at 125pps. Performance with single-channel strategies did not differ significantly across pulse rates above 250pps (p>0.05).

#### 3.2.1. Individual Subjects Results

Individual consonant identification results from 10 participants are shown in Figure 3. The first column presents percent correct scores. The remaining three columns show information transmission of consonant features. Multi-channel and single-channel results are shown by blue squares and green circles, respectively, with error bars representing within-subject 95% confidence intervals. Confidence intervals for correct consonant scores were estimated using binomial distribution (Thornton, Raffin, 1978). For information transfer of consonant features, confidence intervals were calculated using the bootstrapping method (Azadpour et al., 2014).

**Figure 3.**
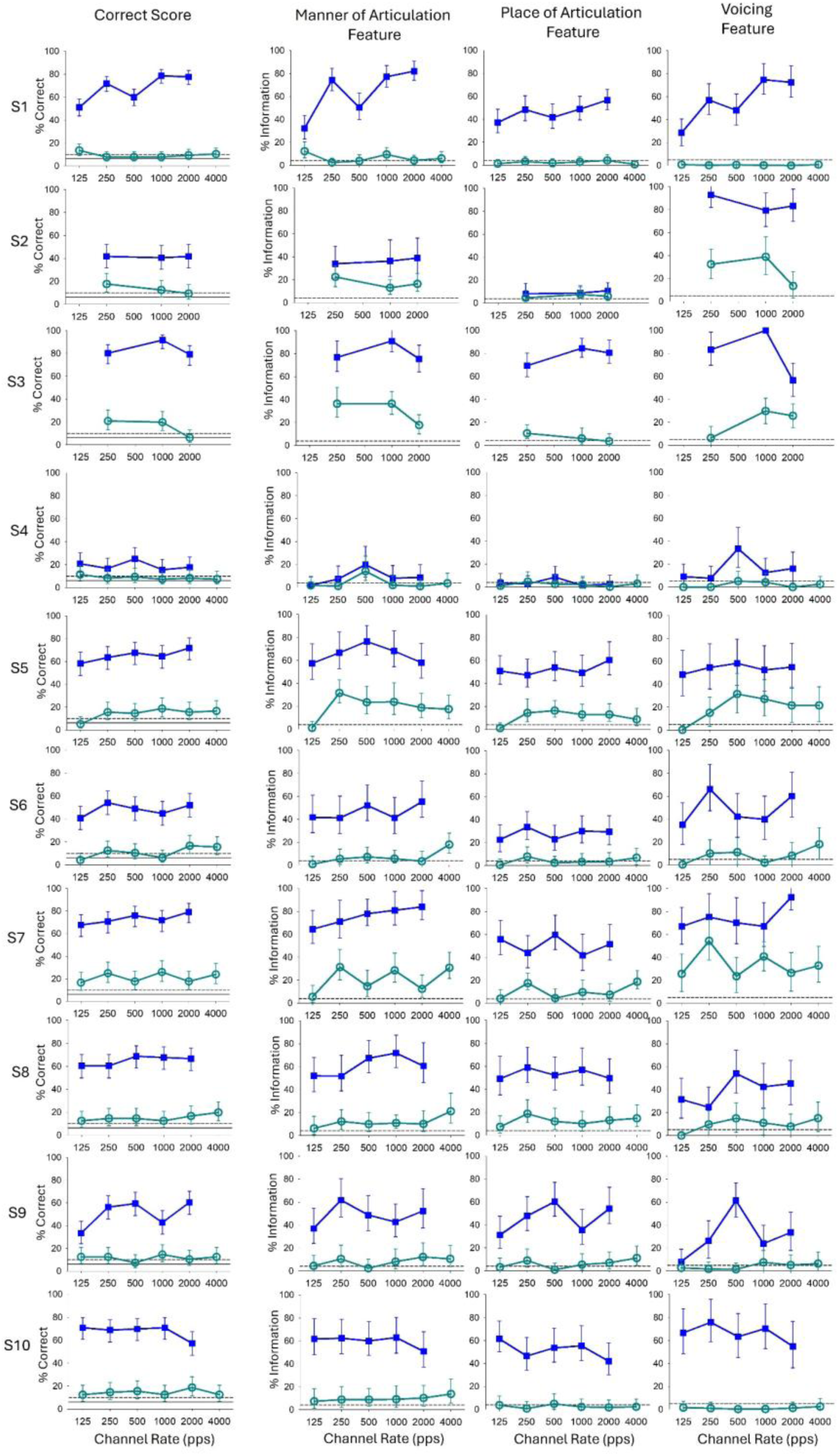
Individual subject results showing the effects of pulse rate on consonant scores (first column) and information transfer of articulation features (columns 2-4). Error bars represent 95% confidence intervals.

Within-subject differences were assessed using the 95% confidence intervals denoted by error bars in Figure 3. Significant differences between strategy conditions within subjects were identified where error bars did not overlap. This analysis revealed significant effects of pulse rate on correct consonant scores in subjects S1, S3, and S9. For S1 and S9, the multi-channel consonant score was significantly lower at 125pps compared to at least one other rate. In S3, the single-channel consonant score was significantly lower at 2000pps, the highest tested rate. Pulse rate also significantly affected information transmission of consonant features in subjects S1, S3, S4, S5, S6, S7, and S9, for multi-channel or single-channel strategies. The effects of pulse rate could vary across features and between single-channel and multi-channel programs. For example, in S1, transmission of manner and voicing features was poorest at 125pps with the multi-channel strategy, whereas manner information transmission at 125pps was the highest for single-channel strategy. In S3, the lowest voicing feature transmission was at 2000pps for the multi-channel strategy and at 125pps for the single-channel strategy. Also, for the single-channel strategy in S3, the best rate for manner of articulation (250pps) was the worst rate for voicing. For S7, manner feature transmission was significantly highest at 4000pps for single-channel strategy, showing benefit from high pulse rates. For S9, voicing feature transmission was highest at 500pps for multi-channel strategy, showing advantage for mid-range pulse rates. These results align with previous findings that pulse rate effects can vary significantly among CI users. Furthermore, the current findings suggest that pulse rate can differentially affect the single-channel and multi-channel transmission of consonant articulation features within individuals.

### 3.3. Amplitude Modulation (AM) Detection

The average AM detection thresholds are presented in Figure 4 for single-channel strategy (green circles) and multi-channel ACE (blue squares) across different pulse rates. Pulse rate effects on AM detection were analyzed using separate one-way repeated measures ANOVA for single-channel and multi-channel results. The analysis revealed significant pulse rate effects for both multi-channel (F(4, 36)=4.35, p=0.006) and single-channel (F(4, 45)=6.38, p<0.001) strategies. Post-hoc Holm-Sidak tests indicated that poorer AM detection at 125pps drove these effects. For multi-channel strategies, AM detection thresholds at 125pps were significantly worse than that of 500, 1000, and 2000pps (p<0.05). Similarly, for single-channel strategies, AM thresholds at 125pps were significantly poorer compared to 250, 500, 1000, and 2000pps (p<0.05), but not compared to 4000pps (p=0.27). No significant differences in AM detection thresholds were observed between 4000pps and other rates (p>0.16). Although the decline at 4000pps for single-channel strategies was not statistically significant, the inverse U-shape pattern in Figure 4 is nonetheless consistent with the hypothesis that elevated perceptual variability at high pulse rates may impair modulation detection, and warrants investigation with larger samples. At 125pps, the decline in AM detection is likely due to psychophysical limitations in temporal processing rather than inadequate AM presentation. The 125pps pulse rate is five times larger than the 25Hz AM frequency, satisfying criteria for accurate envelope sampling (McKay et al., 1994).

**Figure 4.**
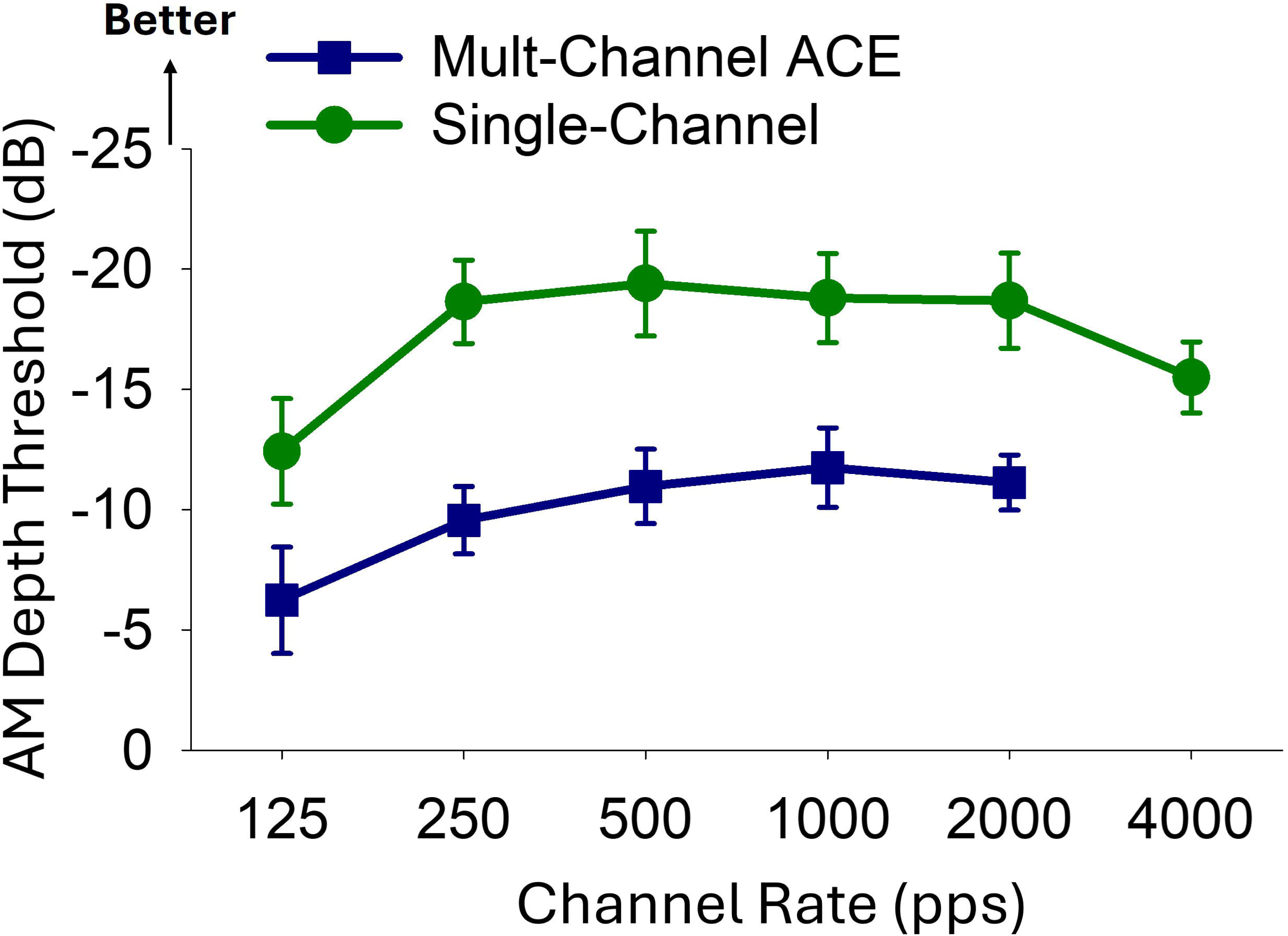
Average AM depth thresholds (in dB) for single-channel (green circles) and multi-channel clinical (blue squares) strategies across pulse rates.

#### 3.3.1. AM detection thresholds for single-channel vs. multi-channel strategies

Figure 4 shows that AM detection performance with multi-channel strategies is considerably poorer than with single-channel strategies. This is counterintuitive as it is expected that AM detection would be more accurate when AM is presented at multiple electrode sites across the array as opposed to a single electrode. To investigate the reasons for this unexpected outcome, we analyzed electrodogram outputs of AM and non-AM noise bands. Figure 5 illustrates electrode current level outputs for multi-channel ACE (top) and single-channel (bottom) strategies, with different colors in the multi-channel figures indicating separate electrodes. T and C levels were set at 100 and 200 CL for these demonstrations. Two key differences between the outputs of multi-channel and single-channel strategies were observed. First, the multi-channel strategy exhibited greater fluctuations in current levels for both AM and non-AM stimuli, likely caused by inherent envelope fluctuations for narrow-band frequency channels. In contrast, the single-channel strategy produced cleaner, less fluctuating current outputs, reflecting the envelope of a broad-band channel. The greater current amplitude fluctuations may contribute to reduced ability to differentiate AM from non-AM stimuli in the multi-channel condition. The second difference observed between the outputs of single-channel and multi-channel strategies was that the multi-channel strategy produced smaller electrode current amplitudes compared to the single-channel strategy. This is presumably due to reduced signal energy within narrow-band channels compared to the broad-band channel used in the single-channel strategy. The lower current amplitude for multi-channel strategy may contribute to poor AM detection, as detection of electric AM at a single-electrode deteriorates at lower stimulation levels (Fraser, McKay, 2012; Galvin, Fu, 2009).

**Figure 5.**
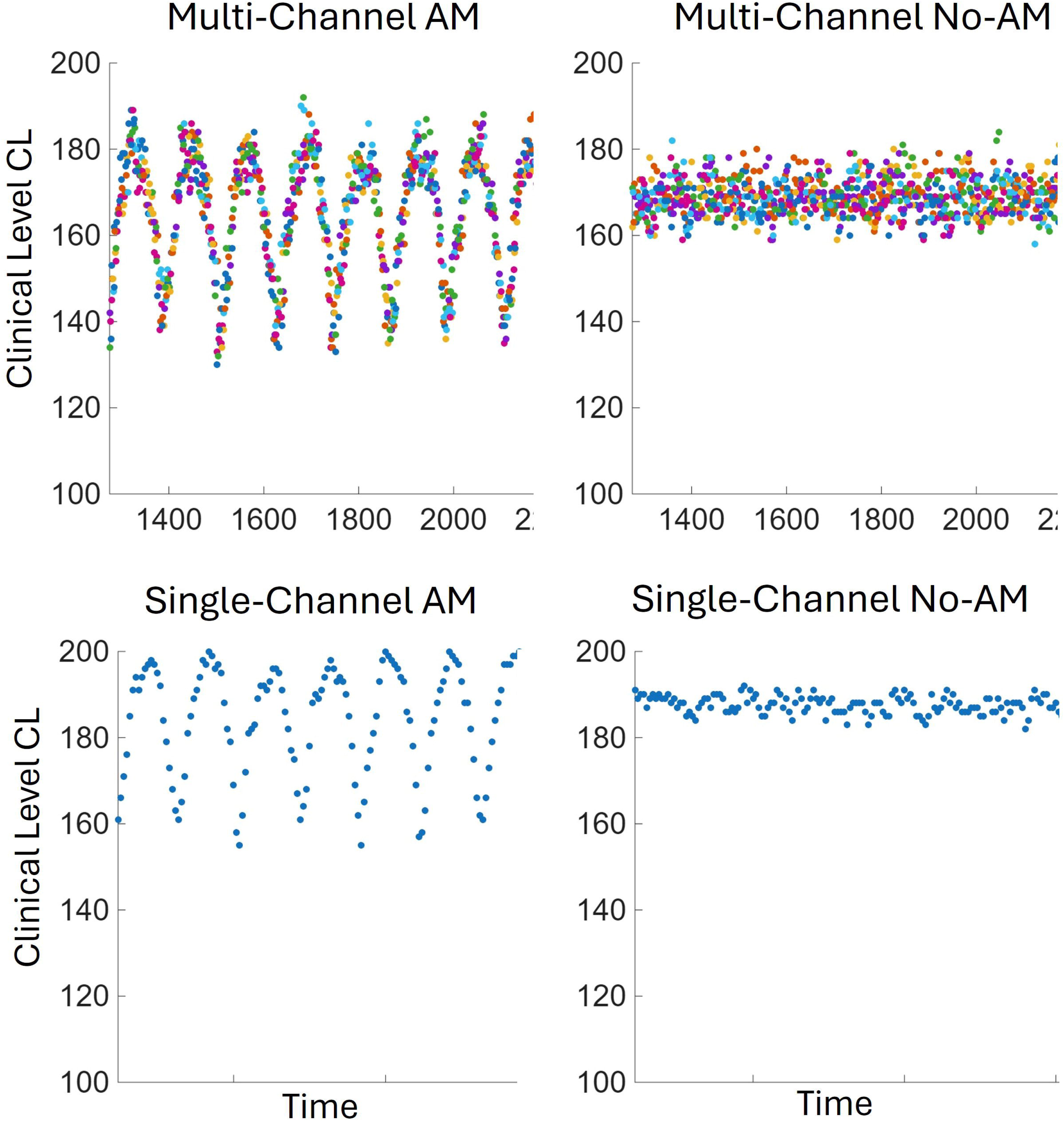
Representative electrode current outputs for amplitude-modulated (AM) and unmodulated (No-AM) stimuli delivered with multi-channel and single-channel stimulation strategies. Different colors indicate current activity on individual electrodes. T and C levels were set at 100 and 200CL for illustration purposes.

### 3.4. Contribution of within-channel temporal processing to multi-channel performance

Outcomes with single-channel CI strategies can provide insights into the transmission of temporal envelope cues at specific cochlear regions, without the influence of cross-channel interactions. We evaluated the role of within-electrode temporal transmission in multi-channel outcomes by evaluating the correlation between single-channel and multi-channel results. The correlation between single-channel and multi-channel performance was evaluated for AM detection and consonant recognition tests with data being averaged across the tested pulse rates. Averaging across pulse rates provided a more reliable composite estimate for each measure and allowed us to perform a single correlation for each test as opposed to multiple correlations for each pair of rates.

Figure 6A illustrates the relationship between AM detection thresholds for single-channel and multi-channel strategies across 10 subjects. Pearson correlation analysis revealed a strong relationship between single-channel and multi-channel AM detection performance (r=0.89, n=10, p<0.001; 95% confidence interval: [0.59, 0.97]). These findings suggest that detection of AM presented to a single cochlear site in the middle of the CI electrode array predicts detection of multi-channel AM presented through clinical strategies.

**Figure 6.**
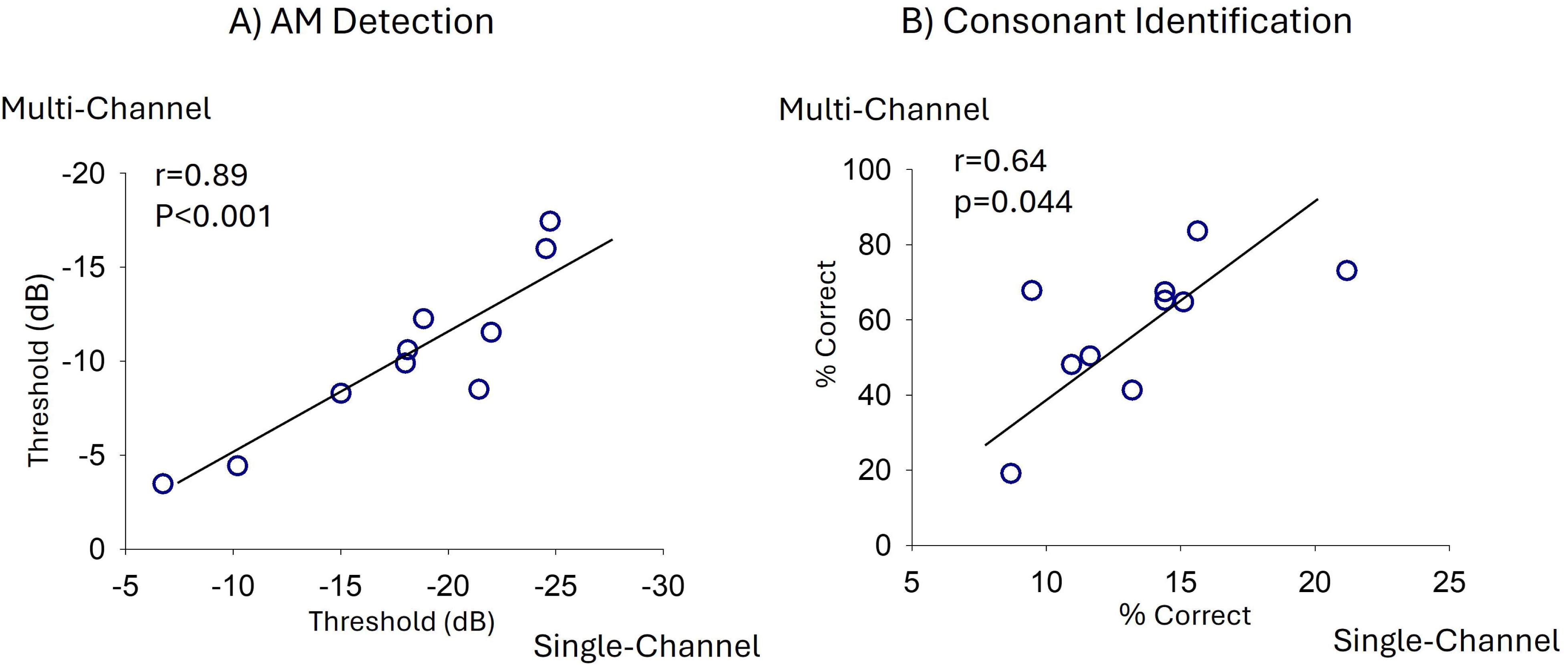
Correlation between multi-channel and single-channel stimulation strategies for AM detection thresholds and consonant identification performance across individual subjects.

Figure 6B illustrates the relationship between correct consonant scores for single-channel and multi-channel strategies, with scores averaged across all tested channel rates. Pearson correlation analysis showed a significant relationship between single-channel and multi-channel consonant identification results (r=0.64, n=10, p=0.044; 95% confidence interval: [0.017, 0.9]), indicating the role of within-channel speech envelope transmission at individual electrodes in supporting broader multi-channel speech recognition performance. The variability in temporal envelope transmission via a single electrode in the middle of the CI array accounted for approximately 40% of the variability in multi-channel consonant recognition outcome.

A marginally significant correlation was also found between multi-channel consonant scores and single-channel transmission of manner of articulation (r=0.63, n=10, p=0.049; 95% confidence interval: [0.0006, 0.9]). However, no significant correlation was found between multi-channel scores and single-channel voicing feature (r=0.29, n=10, p=0.41; 95% confidence interval: [-0.4, 0.8]). The two correlations were marginally but not statistically different from each other (p=0.053), considering the high correlation between manner of articulation and voicing features (r=0.78, n=10, p=0.008). Information transfer for place of articulation was excluded from this analysis as place cues are not reliably represented through single-channel strategies.

### 3.5. Relation between AM detection and consonant identification

Figure 7 shows consonant identification scores plotted against AM detection thresholds for multi-channel (panel A) and single-channel (panel B) strategies, with both metrics averaged across tested pulse rates. Pearson correlation analysis revealed no significant relationship between AM detection thresholds and consonant identification scores for either multi-channel (r=0.25, n=10, p=0.48; 95% confidence interval: [-0.5, 0.8]) or single-channel strategies (r=0.41, n=10, p=0.23; 95% confidence interval: [-0.3, 0.8]). Given the modest sample size (n = 10), these analyses had limited statistical power to detect moderate correlations. The absence of a significant association should therefore be interpreted cautiously; however, the observed correlation coefficients were small to moderate in magnitude, suggesting that 25 Hz AM detection may not be strongly predictive of consonant recognition performance in this paradigm.

**Figure 7.**
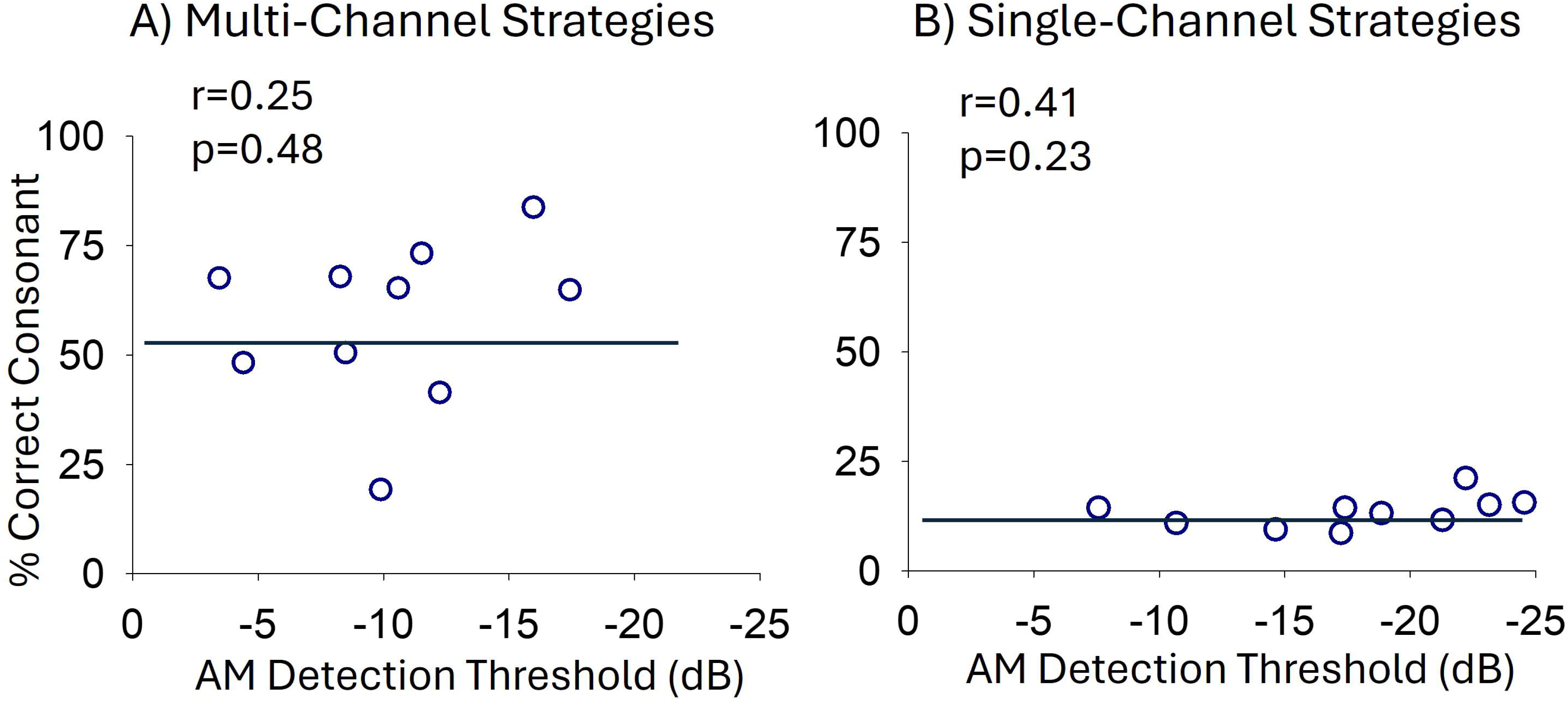
Correlation between consonant recognition performance and AM detection thresholds for multi-channel and single-channel stimulation strategies across individual subjects.

## 4. Discussion

### 4.1. Effect of pulse rate and electrode location on T and C levels

The average T and C levels across the electrode array decreased as pulse rate increased, with T levels showing a larger and more systematic reduction compared to C levels, consistent with previous findings (Kreft et al., 2004; Zhou et al., 2012). The pronounced impact of pulse rate on T levels may be explained by the facilitative effects of pulse interactions at lower current amplitudes (Boulet et al., 2016; Middlebrooks, 2004). Analysis of T and C levels across the array revealed significantly higher values for mid-array electrodes compared to apical electrodes at low pulse rates (125pps and 250pps). This observation is consistent with previous reports on T level patterns obtained with perimodiolar electrode array designs (Gordon, Papsin, 2013) and may reflect the larger electrode-to-modiolus distances in the mid-cochlear region for these electrode arrays (Long et al., 2014). Notably, the across-array differences in T and C levels diminished at higher pulse rates. It has been proposed that T levels provide insights into the local electrode-neural interface quality at each electrode site (Bierer, 2010; Long et al., 2014; Pfingst, Xu, 2004). The observed reduction in spatial variations in T levels at higher stimulation rates may result from temporal pulse interactions broadening the spatial spread of neural excitation and reducing local specificity (Middlebrooks, 2004; Zhou, 2016). This suggests that low-rate stimulation may be more effective for identifying local electrode-neural interface quality across CI arrays.

### 4.2. Effect of pulse rate on consonant recognition

Consonant recognition scores were similar across mid to high pulse rates between 250 and 2000pps. This range includes the lowest and highest pulse rates that are typically used in clinical CI strategies. However, consonant recognition significantly declined at 125pps for both single-channel and multi-channel ACE strategies, demonstrating a consistent pattern of reduced speech perception at low pulse rates. This pattern was also observed for information transmission of consonant features, where transmission of manner of articulation feature was consistently poorer for 125pps compared to higher rates. The decline at 125pps is consistent with perceptual constrains on temporal envelope processing at low carrier rates, rather than inadequate sampling of the acoustic envelope. Critical speech envelope variations are predominantly below 20Hz (Sinha, Azadpour, 2024; Souza, Rosen, 2009), and should be precisely sampled at 125pps carrier rate (McKay et al., 1994). The poorer performance at 125pps likely reflects perceptual inefficiencies at lower pulse rates rather than encoding limitations.

### 4.3. Variability in pulse rate effects across individuals

Individual subject results revealed significant variability in the effects of pulse rate on consonant recognition and information transfer of articulation features. This observation is consistent with previous indications that optimal pulse rates can vary across individuals (Plant et al., 2007; Shader et al., 2020; Weber et al., 2007). The results of this study show that pulse rate can differentially affect the transmission of consonant articulation features across individuals, meaning that optimal pulse rate may be different for different consonant features within an individual. This may explain the overall flat pattern of consonant recognition scores as a function of pulse rate. Additionally, pulse rate can affect single-channel and multi-channel outcomes differently. The inconsistency between single-channel and multi-channel strategies reveals that pulse rate influences within-channel and cross-channel speech cues differently. These observations suggest that pulse rate benefits depend on individual auditory profiles that determine the transmission of temporal and spectral cues.

### 4.4. AM detection and partial support for inverse U-shaped pattern

AM detection results were partially consistent with an inverse U-shaped function of pulse rate. The decline at 125 pps was statistically robust, while the decline at 4000 pps, though descriptively present, did not reach significance. These results provide partial support for the hypothesis that very high pulse rates impair modulation sensitivity, but larger samples with greater statistical power are needed to confirm the high-rate effect. The poorer AM detection at 125pps is likely due to psychophysical limitations in modulation sensitivity rather than imprecise envelope sampling, as the 25Hz modulation frequency used in this study meets the minimum criteria for accurate sampling at 125pps (McKay et al., 1994). Notably, poorer AM detection at 125pps is consistent with poorer consonant recognition at this rate, suggesting that reduced modulation transmission at low pulse rates may underlie poorer speech perception at these rates. The non-significant reduction in single-channel AM detection performance at 4000pps aligns with previous findings that high pulse rates impair sensitivity to amplitude variations (Galvin, Fu, 2005; Galvin, Fu, 2009), and may result from increased loudness variability (Azadpour et al., 2018). However, the decline at 4000pps was not statistically significant, suggesting that perceptual effects at high rates may be inconsistent and subject to individual variability.

AM detection performance with single-channel strategies was significantly better than with multi-channel strategies. This is a counterintuitive observation given the expectation that multi-channel stimulation across the cochlea would enhance the representation of amplitude modulations. Electrodogram analyses revealed that multi-channel strategies result in greater fluctuations in current amplitudes and lower overall stimulation levels compared to single-channel strategies. Envelope fluctuations in narrow band channels and reduced channel-specific current levels may explain the poorer AM detection performance observed for multi-channel approaches (Fraser, McKay, 2012; Galvin, Fu, 2009).

### 4.5. Role of single-channel temporal envelope in multi-channel outcomes

The correlations between single-channel and multi-channel performance emphasize the importance of temporal envelopes at individual cochlear sites in supporting broader multi-channel outcomes. AM detection thresholds with single-channel strategies strongly predicted AM thresholds with multi-channel strategies (r=0.89), demonstrating that temporal cue transmission at a single electrode plays a critical role in multi-channel responses. Similarly, single-channel speech recognition performance moderately correlated with multi-channel outcomes (r=0.64), with variability in single-electrode envelope transmission accounting for approximately 40% of the variance in multi-channel speech recognition results.

These findings support an important contribution of within-channel temporal envelope processing to multi-channel speech perception outcomes in cochlear implant users. Single-channel measures provide a valuable approach for evaluating envelope transmission at individual cochlear sites, allowing researchers and clinicians to isolate and assess temporal processing capabilities independent of cross-channel spectral representations. Single-channel speech assessments used in this study may offer a means of identifying electrode-specific contributions to speech cue transmission in CIs.

### 4.6. Absence of significant correlation between AM detection and consonant recognition

In contrast to some previous studies (Fu, 2002; Won et al., 2011), this study didn’t find a significant correlation between AM detection thresholds and consonant identification scores for either single-channel or multi-channel strategies. This absence of correlation may be attributable to the relatively low modulation frequency of 25 Hz used for AM stimuli in the present study, compared to 100Hz or larger modulation frequencies used by Fu (2002) and Won et all (2011). Although 25 Hz falls within the range of speech-critical envelope frequencies (Sinha, Azadpour, 2024; Souza, Rosen, 2009), these findings show that AM detection at speech-critical modulation frequencies does not strongly predict speech perception performance, suggesting limited clinical utility for assessing speech outcomes. The reasons why significant correlations have been reported at higher modulation frequencies, beyond those considered critical for speech envelope encoding, warrants further investigation.

### 4.7. Limitations and future directions

This study has several limitations that should be considered. The modest sample size (n = 10) consisted exclusively of post-lingually deafened adults using Cochlear Nucleus devices with perimodiolar electrode arrays, which limits the generalizability of findings to other CI systems and patient populations. The multi-channel experimental strategies employed a fixed maxima of 6, differing from the standard clinical configurations. Additionally, single-channel testing was conducted using a single mid-array electrode (electrode 12), and the findings may not generalize to apical or basal regions of the cochlea. Only one AM frequency (25 Hz) was tested; evaluating a broader range of modulation frequencies could provide a more comprehensive understanding of temporal processing across pulse rates. While T and C levels were measured at each rate to control for rate-dependent loudness differences, residual confounds related to rate-dependent changes in loudness growth cannot be entirely ruled out.

Despite these limitations, the findings of this study highlight the complex relationship between pulse rate, temporal envelope processing, and consonant recognition in CI users. Mid-range pulse rates generally produced consistent performance, but the variability observed across individuals underscores the importance of personalized programming strategies to optimize CI outcomes. Single-channel assessments of modulation detection and consonant identification provide valuable insights into the transmission of temporal cues at specific cochlear sites. Future research should conduct more detailed analyses of within-channel and cross-channel envelope representations to better understand the factors driving speech perception variability. Such efforts can guide the development of personalized approaches to enhance speech outcomes, particularly for CI users with poor performance.

## Declaration of generative AI and AI-assisted technologies in the manuscript preparation process

During the preparation of this work, the authors used ChatGPT to improve readability and clarity. The authors reviewed and edited the output as needed and take full responsibility for the content of the published article.

